# scRNA-seq analysis of colon and esophageal tumors uncovers abundant microbial reads in myeloid cells undergoing proinflammatory transcriptional alterations

**DOI:** 10.1101/2020.05.14.096230

**Authors:** Welles Robinson, Joshua K. Stone, Fiorella Schischlik, Billel Gasmi, Michael C. Kelly, Charlie Seibert, Kimia Dadkhah, E. Michael Gertz, Joo Sang Lee, Kaiyuan Zhu, Lichun Ma, Xin Wei Wang, S. Cenk Sahinalp, Rob Patro, Mark D.M. Leiserson, Curtis C. Harris, Alejandro A. Schäffer, Eytan Ruppin

## Abstract

The study of the tumor microbiome has been garnering increased attention. We developed a computational pipeline (CSI-Microbes) for identifying microbial reads from single-cell RNA sequencing (scRNA-seq) data. Using a series of controlled experiments and analyses, we performed the first systematic evaluation of the efficacy of recovering microbial UMIs by multiple scRNA-seq technologies, which identified the newer 10x chemistries (3’ v3 and 5’) as the best suited approach. Based on these findings, we analyzed patient esophageal and colorectal carcinomas and found that reads from distinct genera tend to co-occur in the same host cells, testifying to possible intracellular polymicrobial interactions. Microbial reads are disproportionately abundant within myeloid cells that upregulate proinflammatory cytokines like *IL1Β* and *CXCL8* and downregulate antigen processing and presentation (APP) pathways. The latter, however, are markedly upregulated in infected tumor cells. These results testify that intracellular bacteria predominately reside within co-opted myeloid cells, which inflame the tumor microenvironment and may influence immunotherapy response.

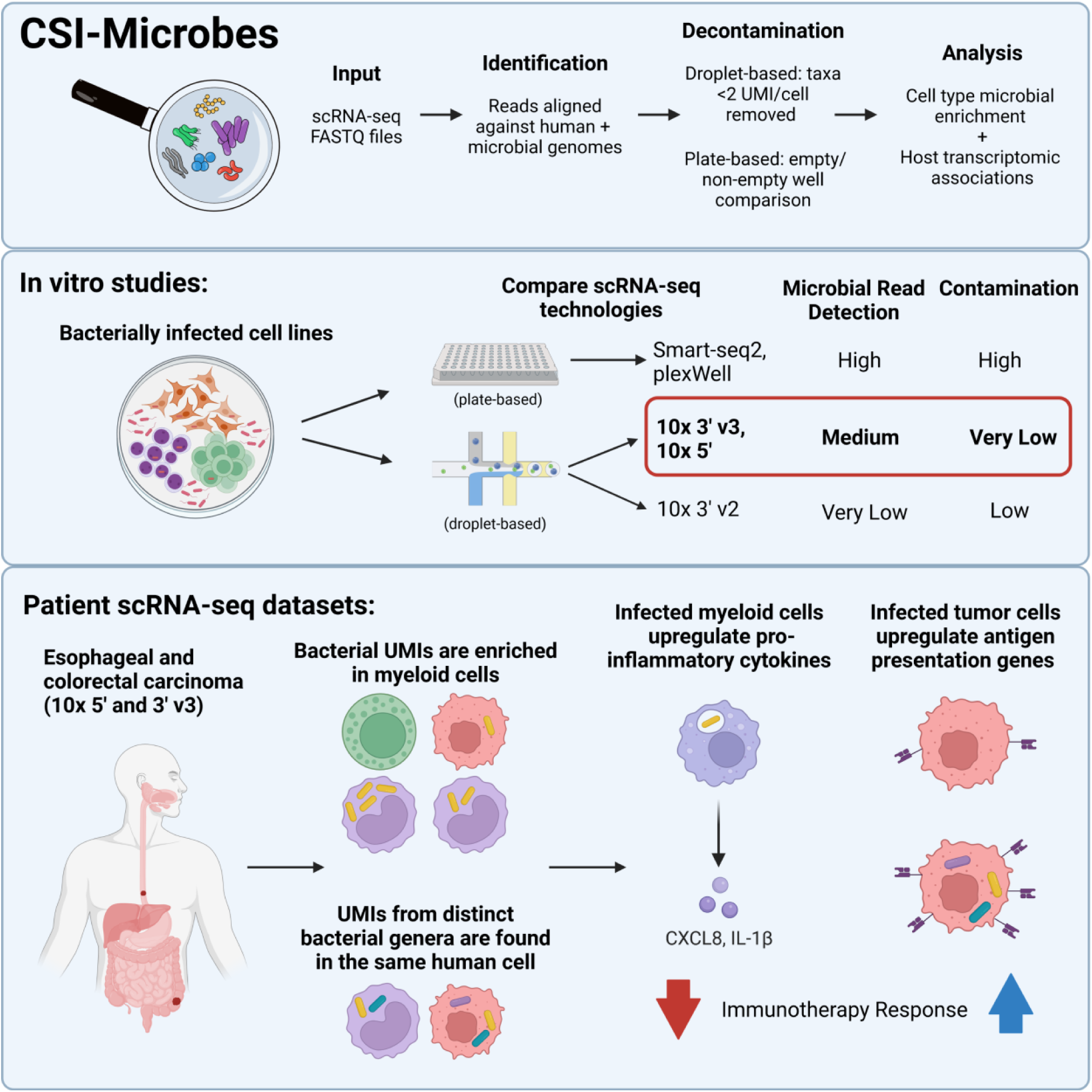

## Introduction

In addition to malignant and non-malignant human cells, the tumor microenvironment (TME) consists of microbes including viruses, bacteria, and fungi, collectively referred to as the tumor microbiome. Early studies of the tumor microbiome focused on viruses that are estimated to cause ∼10-15% of human cancers worldwide, including *Merkel polyomavirus*, which is detectable in ∼75% of Merkel cell carcinomas and *Hepatitis B* and *C viruses*, which are collectively estimated to cause more than 60% of liver cancers^1–4^.

Some more recent experimental and computational studies expanded the scope of the tumor microbiome to include tumor-resident bacteria and fungi^5–8^. For example, early studies of bacteria in tumors reported the increased prevalence of the bacterium *Fusobacterium nucleatum* in colorectal carcinoma compared to adjacent non-tumor tissue^9, 10^. Larger-scale reports demonstrate that many, if not all, solid tumor types have a microbiome, possibly distinct and distinguishable from the microbiome of nearby non-tumor tissue^6, 8–10^. Further studies have demonstrated the functional importance of specific members of the tumor microbiome to multiple hallmarks of tumorigenesis including mutagenesis, metastasis, and immune evasion as well as response to chemotherapy^9–15^. Tumor microbiome studies have shifted partly to intracellular microbes due to recent findings that the vast majority of intratumoral bacterial taxa, including *F. nucleatum,* appear to reside intracellularly within the tumor microenvironment^6, 7, 16^. Despite these advances, it has remained an important, open challenge to identify which microbial taxa reside intracellularly and whether they reside exclusively or preferentially inside tumor cells, immune cells, or cells of the non-cancerous tissue adjacent to the solid tumor.

One increasingly popular approach for studying intracellular or host cell-associated microbes is the analysis of microbial reads from single-cell RNA sequencing (scRNA-seq) in the context of viral infection or of cancer^17–21^. To provide context for our work, we mention two recently published reports analyzing microbial reads from scRNA-seq in the context of cancer^19, 21^. One of these studies introduced a novel computational approach (SAHMI) for decontaminating droplet-based scRNA-seq data, which it applied to analyze two 10x 3’ v2 datasets of pancreatic cancer^19^. The other introduced a novel experimental scRNA-seq approach and analysis pipeline termed INVADEseq that aims to increase the capture of bacterial reads by including a primer for a conserved region of the bacterial 16S rRNA in 10x 5’ polyA capture, which was applied to analyze seven oral squamous cell carcinomas (OSCC)^21^. These studies present distinct methods for filtering out potential environmental contaminants^22^, which has been one of the primary concerns with the application of traditional 16S rRNA-seq and bulk DNA- and RNA-seq-based microbiome studies. SAHMI uses multiple decontamination steps while the INVADEseq study does not discuss methods for filtering potential contaminants. While making important contributions, each of these two studies focused on studying one single scRNA-seq wet lab method, leaving open the challenge of providing comparative guidance regarding scRNA-seq technology selection for future scRNA-seq studies that profile the microbiome.

To compare microbial read detection of different scRNA-seq technologies, we developed a reproducible computational pipeline, named CSI-Microbes. We then comprehensively applied CSI-Microbes to study multiple plate-based (Smart-seq2 and plex-Well) and droplet-based (10x 3’ v2, 3’ v3, and 5’) scRNA-seq datasets of human cells exposed *in vitro* to select bacteria. Specifically, we found that plate-based technologies capture the most microbial reads but approximately half of these microbial reads map to putative contaminant genera. By comparison, 10x technologies capture relatively few reads from putative contaminants but more successfully capture reads mapping to the *in vitro* exposed bacteria. Their capture levels depend on the specific chemistry, with at least an order of magnitude more microbial reads detected by the newer 10x chemistries (10x 3’ v3 and 10x 5’) compared to the earlier 10x 3’ v2 chemistry. These findings thus identify the newer 10x protocols as the preferred methods for studying the tumor microbiome.

Armed with these insights, we next applied CSI-Microbes to interrogate the intratumoral microbiome of patient colorectal and esophageal carcinomas by analyzing large, recently published 10x 3’ v3 and 5’ datasets. We show that intracellular bacteria are disproportionately found in myeloid cells and have identified bacterial genera that co-occur in the same cells more than expected under a random model. Finally, we combined the microbial and host cells transcriptomic reads to reveal cell type-specific transcriptomic signatures that are shared between *in vitro* and *in vivo* infected cells. Those include the up-regulation of antigen processing and presentation pathways in infected tumor cells and the up-regulation of proinflammatory cytokines *IL1Β* and *CXCL8* in infected myeloid cells, whose potential therapeutic relevance is further discussed.

## Results

### Overview of CSI-Microbes

We developed a computational pipeline, *CSI-Microbes* (https://github.com/ruppinlab/CSI-Microbes-identification), to identify microbial reads from plate-based and 10x scRNA-seq datasets (when comparing plate-based and 10x datasets, we use the word ‘reads’ to refer to unique molecular identifiers (UMIs) when describing 10x data for consistency) (**Figure S1A**). The modules of CSI-Microbes are described in **STAR Methods**; for some of the more technical parts of Results, it is useful to be aware that the module that aligns reads to genomes can use different approaches for aligning non-human reads to microbial genome(s) including PathSeq^23^ (which was used also by the INVADEseq study) and CAMMiQ^24^ to align to many microbial genomes and SRPRISM^25^ to align to a (few) specific microbial genome(s). Unless otherwise specified, the results presented in this study use the PathSeq option for sequence alignment. In total, we applied CSI-Microbes to seven different datasets in this study (**Table 1**).

**Table 1.** Datasets Analyzed.

| Abbreviation | scRNA-seq technology | Description | Accession | Source |
| --- | --- | --- | --- | --- |
| Aulicino2018 | Smart-seq2 (plate-based) | moDCs exposed to *S. enterica* | PRJNA437328 | <sup>26</sup> |
| Ben-Moshe2019 | 10x 3' v2 | PBMCs exposed to *S. enterica* | PRJNA503437 | <sup>27</sup> |
| Paulson2018 | 10x 3' v2 and 5' | Merkel cell carcinoma | PRJNA483959<br>and<br>PRJNA484204 | <sup>33</sup> |
| Robinson2023 | plexWell (plate-based) and 10x 3' v3 and 5' v2 | HCT116, THP1 and Jurkat T exposed to *F. nucleatum* |  | This paper |
| Pelka2021 | 10x 3' v2 and 3' v3 | Colorectal carcinoma | PRJNA723926 | <sup>37</sup> |
| Ma2019 | 10x 3' v2 and 3' v3 | Hepatocellular carcinoma | PRJNA636285 | <sup>31,32</sup> |
| Zhang2021 | 10x 5' | Esophageal carcinoma | PRJNA672851 | <sup>36</sup> |

The Results section is composed of two main parts, A and B, each composed of a few subsections. **Part A** describes a series of controlled experiments and analyses that evaluate the efficacy of different scRNA-seq technologies in uncovering the microbiome of infected cells *in vitro*. Based on these findings, **Part B** transitions to analyze patients’ tumor data using the most informative sequencing platforms identified in part A to interrogate host-microbe interactions.

### A. Evaluating CSI-Microbes across *in vitro* experiments and sequencing platforms

#### CSI-Microbes identifies reads from known intracellular microbes from specifically designed plate and droplet-based scRNA-seq technologies

To test and validate our pipeline, we first analyzed two publicly available scRNA-seq datasets of human cells exposed *in vitro* to the known invasive bacterium *Salmonella enterica* in a controlled manner. Aulicino2018 is a plate-based scRNA-seq (Smart-seq2) dataset of monocyte-derived dendritic cells (moDCs) exposed to *S. enterica* as well as unexposed control cells^26^. The authors divided the exposed moDCs into “infected” or “bystander” cells depending on whether the presence of CellTrace-labeled *S. enterica* could be detected using FACS. Ben-Moshe2019 is a droplet-based (10x 3’ v2) scRNA-seq dataset of human peripheral blood mononuclear cells (PBMCs) exposed to *S. enterica* as well as unexposed control cells^27^. These authors used FACS to reveal that monocytes were enriched for the red fluorescent protein (RFP) expressed by *S. enterica* in their model system.

Analysis of these two datasets with CSI-Microbes identified a very wide range in the number reads mapping to the *Salmonella* genus from 756,327 reads in Auclino2018 (mean 2,886 reads per cell) to 8 reads from Ben-Moshe2019 (mean 0.0023 reads per cell). *Salmonella* reads were enriched in the “infected” cells in Aulicino2018 and in monocytes in Ben-Moshe2019, comparable with the author’s FACS results^26, 27^ (**Figures 1A, 1B**). These studies were not designed for the analysis of microbial reads and share one important limitation when adapted for this purpose: the use of primary human blood cells *ex vivo*, which could have been exposed to other microbes *in vivo* that may still be detectable by sequencing. Therefore, it is not possible to determine whether reads that map to genera other than *Salmonella* represent contaminants or prior exposure, which confounds comparing the number of reads from (potentially) contaminating microbes captured by each technology.

**Figure 1:**
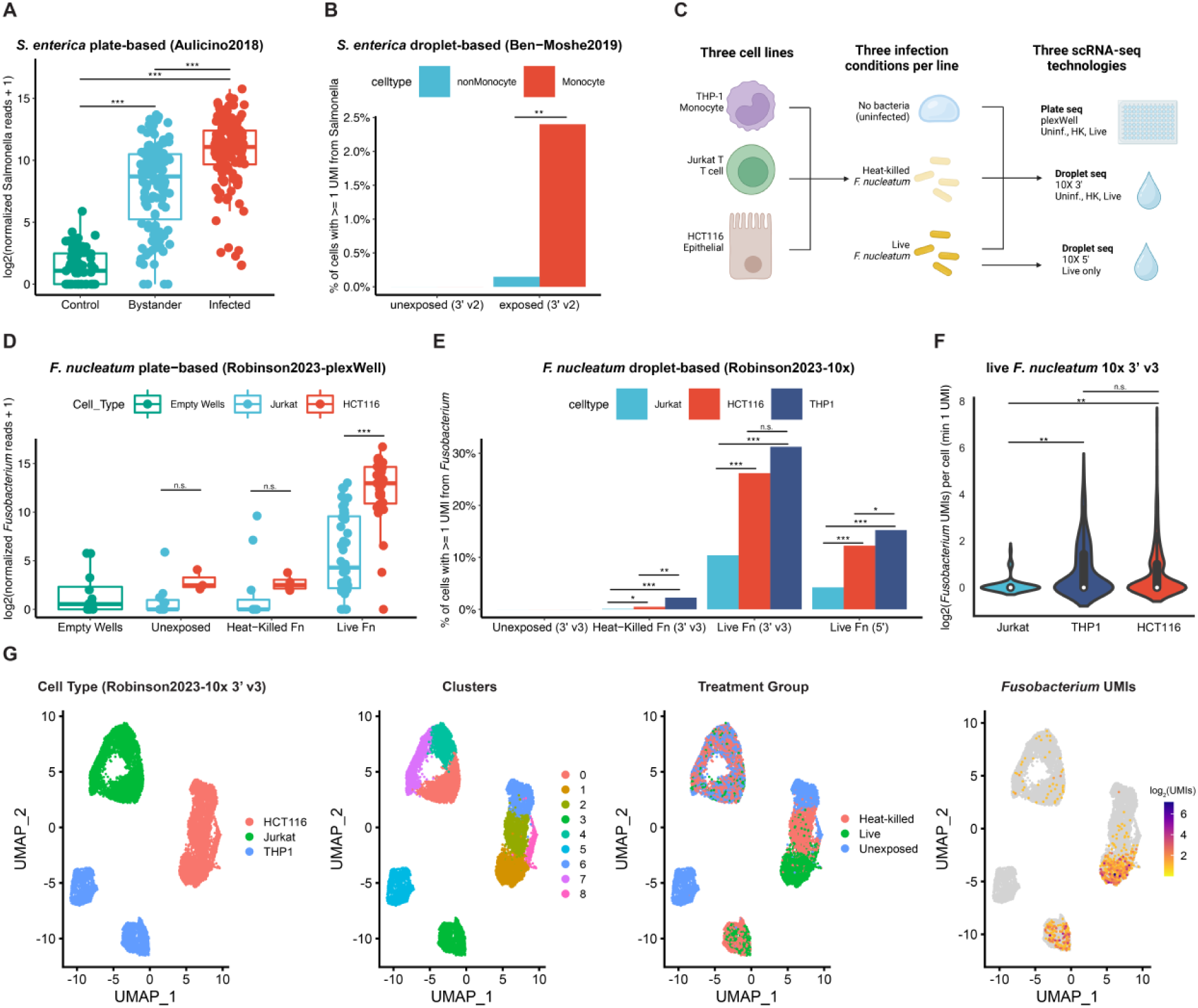
Comparison of the performance of CSI-Microbes on droplet vs plate-seq platforms. **(A)** The number of reads (spike-in normalized and log_2_ transformed) mapping to Salmonella per monocyte-derived dendritic cell (moDC) grouped by exposure condition and sequenced using plate-based scRNA-seq by Aulicino2018. **(B)** The percentage of PBMC cells with at least one read from Salmonella grouped by cell type and exposure condition sequenced using 10x 3’ v2 by Ben-Moshe2019. **(C)** Experimental design for F. nucleatum exposure and scRNA-seq for this paper (Robinson2023). **(D)** The number of reads (spike-in normalized and log2 transformed) per cell mapping to Fusobacterium from Jurkat T and HCT116 cells grouped by cell type and exposure condition and sequenced using plate-based scRNA-seq (Robinson2023-plexWell). **(E)** The percentage of cells with at least one read from Fusobacterium from Jurkat T, HCT116 and THP1 cells grouped by cell type and exposure condition sequenced using 10x 3’ v3 and 5’ (Robinson2023). **(F)** The number of UMIs from Fusobacterium per cell grouped by cell type (only cells with ≥ 1 Fusobacterium UMI were included) **(G)** UMAP visualization of cells sequenced using 10x 3’ v3 including cell type, transcriptomic cluster, treatment group and Fusobacterium UMIs. * p-value < 0.05; ** p-value < 0.01; *** p-value < 0.001.

To compare the capture of bacterial reads by different scRNA-seq technologies, we designed new *in vitro* infection experiments using additional controls, which we assayed using multiple scRNA-seq technologies (**Figure 1C**). First, we selected three scRNA-seq technologies to compare using the same experimental design: 10x 3’ v3 and 10x 5’ (droplet) and plexWell (plate). Second, we included three cell lines of various tissue backgrounds, including epithelial cells (HCT116, colorectal cancer), monocytes (THP1) and T cells (Jurkat T). Third, we utilized three experimental treatment groups for each cell line: naïve or unexposed cells for background microbial UMI signal, heat-killed *F. nucleatum*, and live *F. nucleatum* exposed cells. Fourth, we included empty wells in the plexWell capture to account for well carryover as a possible source of contamination.

Analysis of the number of reads mapping to *Fusobacterium* from this study (which we term Robinson2023, and its technology-specific subsets as Robinson2023-10x and Robinson2023-plexWell) was consistent with our prior analysis of the *Salmonella* datasets. We identified 393,380 reads mapping to *Fusobacterium* (mean 4,574 reads per cell) from the plate-based technologies compared to 1,401 reads from 10x 3’ v3 (mean 0.51 reads per cell) and 926 reads from 10x 5’ (mean 0.19 reads per cell), and these differences were significant even after correcting for sequencing depth (Fisher exact test p-value<1e^-300^; odds ratio=232). We next aligned the non-human reads directly to the *F. nucleatum* genome using SRPRISM^25^ to analyze the type of bacterial transcripts identified because bacterial mRNAs are polyadenylated at a much lower frequency and with significantly shorter polyA tails compared to eukaryotic mRNA^28^. We found that most bacterial transcripts captured by scRNA-seq protocols are rRNA (82%, 92% and 95% of plexWell, 10x 5’ and 10x 3’ v3 *F. nucleatum* reads) and not mRNA (**Figure S1C, SI**), which is reassuringly comparable to previous reports from bulk polyA-capture RNA-seq and 10x Visium spatial transcriptomics^21, 29^.

Next, we grouped cells by their cell line and observed an enrichment of *Fusobacterium* reads in HCT116 (and THP1 cells when included) compared to Jurkat T cells exposed to live *F. nucleatum* in both technologies (**Figures 1D, 1E, 1F**). Overlaying *Fusobacterium* reads and treatment groups onto cell clusters (derived using only human genes) revealed that host transcriptomic changes to bacteria were cell line-dependent (and more pronounced in the cell lines enriched with *Fusobacterium* reads) and that the clustering directly corresponded with *Fusobacterium* reads (**Figure 1G, Figure S1B**).

#### Contamination is a significantly bigger problem in plate-based compared to droplet-based technologies

We then expanded our analysis to reads that mapped to other, unexpected microbial (bacterial, fungal, and viral) genera, which likely represent environmental contaminants. We found that 55% (393,380/710,630) of the genera-resolution microbial reads from cells exposed to live *F. nucleatum* mapped to *Fusobacterium* sequenced using plexWell compared to 98% (926/947) and 99% (1,401/1,411) of the microbial reads from 10x 5’ and 3’ v3, respectively (**Figure 2A**). We found 43% (756,327/1,742,908) of the microbial reads mapped to *Salmonella* in Aulicino2018, which is comparable to Robinson2023-plexWell (**Figure 2B**). In contrast, we found only 22% (8/36) of the microbial reads mapped to *Salmonella* from Ben-Moshe2019, which is a significantly lower percentage than Robinson2023-10x and likely biased by the paucity of total microbial reads in this dataset (**Figure 2B**). We also observed contamination in the form of reads from *Fusobacterium* and *Salmonella* in unexposed control cells in plate but not droplet-based technologies (**Figures 1A-1D, SI**), which likely result from experimental carryover within the same plate as both unexposed and exposed to *F. nucleatum* cells were sequenced on the same plate. In summary, our analysis demonstrates that contamination is a bigger problem in plate-based technologies compared to droplet-based technologies, which is consistent with the idea that contaminants are more likely to reach an exposed well handled by people compared to droplets controlled by a microfluidics system.

**Figure 2:**
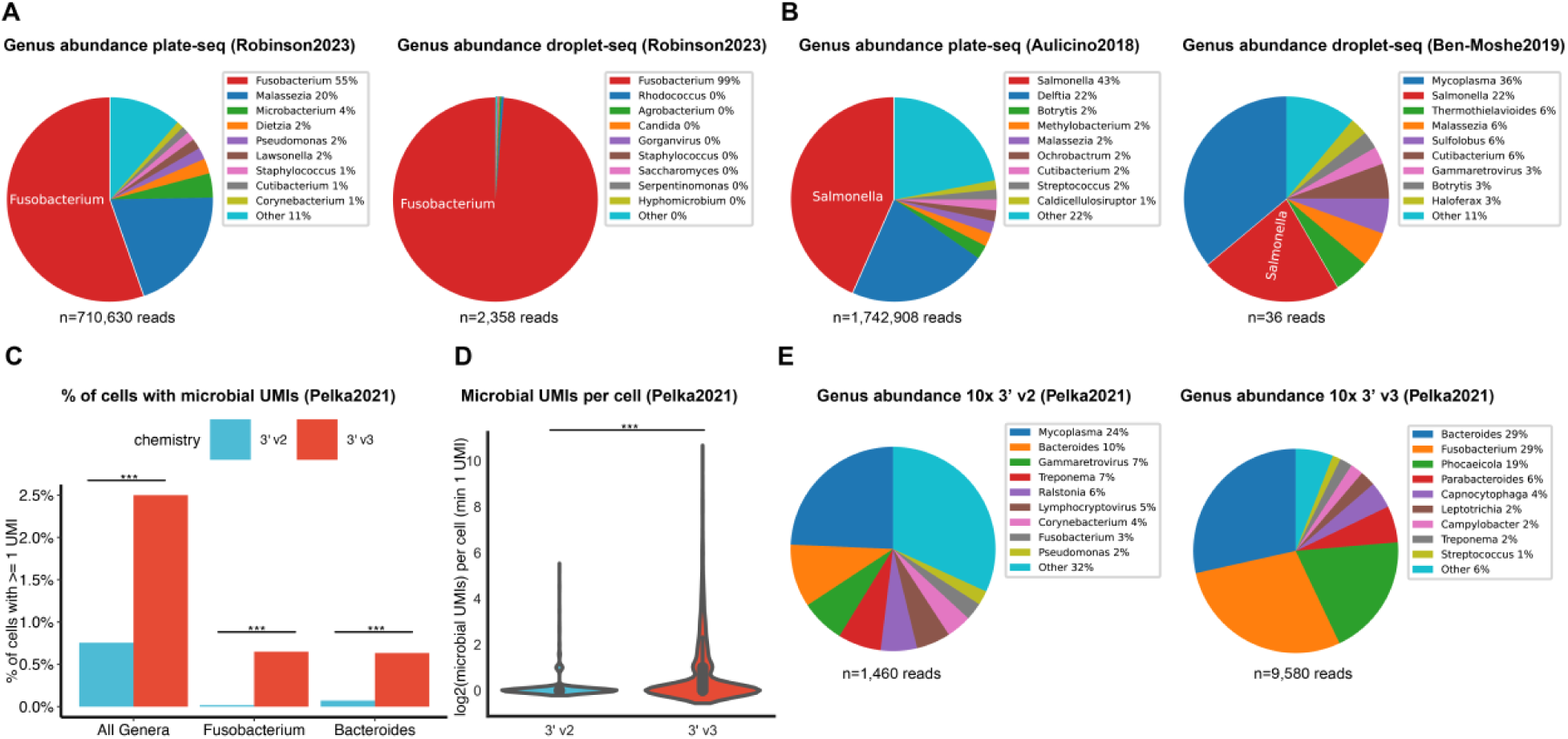
CSI-Microbes reveals numerous contaminant taxa and differences in bacterial reads capture capacities between 10x chemistries. (A) Comparison of the percentage of genera-resolution microbial reads mapped to Fusobacterium and other genera (suspected contaminants) from cells exposed to live F. nucleatum and sequenced using either plate or droplet-based scRNA-seq technologies. (B) Comparison of the percentage of genera-resolution microbial reads mapped to Salmonella and other genera (suspected contaminants) from cells exposed to live S. enterica and sequenced using either plate or droplet-based scRNA-seq technologies. (C) Comparison of the percentage of cells with ≥ 1 microbial UMI between cells sequenced using 3’v2 and 3’ v3 by Pelka2021. (D) Comparison of the number of microbial UMIs per cell (minimum ≥ 1 microbial UMI) between cells sequenced using 3’ v2 and 3’ v3 by Pelka2021. (E) Comparison of the percentage of genera-resolution microbial reads mapped to the top ten genera between 3’ v2 and 3’ v3 chemistries by Pelka2021. * p-value < 0.05; ** p-value < 0.01; *** p-value < 0.001.

#### 10x 3’ v3 and 5’ capture more microbial reads than 10x 3’ v2 chemistries

Although there are important technical differences between the *F. nucleatum* and *S. enterica* invasion experiments, we hypothesized that some of the difference in number of reads captured from the infecting bacterium in the 10x datasets may be related to which chemistry was used, which has been shown to strongly influence which human genes are sequenced (https://kb.10xgenomics.com/hc/en-us/articles/360026501692-Do-we-see-a-difference-in-the-expression-profile-of-3-Single-Cell-v3-chemistry-compared-to-v2-chemistry-). To test for differences in chemistry within a single dataset, we analyzed Pelka2021, which is a large scRNA-seq atlas of colorectal carcinomas sequenced with either 10x 3’ v2 or 10x 3’ v3 chemistries. Even though nearly twice as many cells were sequenced using 10x 3’ v2, we identified substantially more microbial reads from the tumor samples sequenced using 10x 3’ v3 compared to those sequenced using 10x 3’ v2 using the absolute number of microbial reads (9,580 vs. 1,460 genera-resolution microbial reads, respectively), the sequencing depth normalized number (Fisher exact test p-value<1e^-300^; odds ratio=6.24), the percentage of cells with ≥ 1 microbial UMI (Fisher exact test p-value < 4.6e^-260^; odds ratio=3.4) and the number of microbial UMIs per cell (Wilcoxon rank sum test p-value=4e^-30^) (**Figures 2C, 2D**). We also observed differences in the genera detected (**Figure 2E**) and noted that a significantly higher percentage (64% or 6,155/9,580) of the 3’ v3 microbial reads map to genera found enriched in colorectal carcinoma by a meta-analysis study^30^ compared to only 23% (342/1,467) of the 3’ v2 microbial reads (hypergeometric enrichment test p-value < 3e^-194^). The increased number of microbial UMIs combined with the higher proportion of these UMIs mapping to known resident microbes provides an estimate that 10x 3’ v3 captures 36-fold more resident microbial reads per cell compared to 10x 3’ v2. We observed similar results comparing both 10x 3’ v3 and 5’ to 3’ v2 in the analysis of viral reads for *Hepatitis B* in liver cancer (Ma2019^31, 32^) and *Merkel polyomavirus* in Merkel cell carcinoma (Paulson2018^33^) (**Figures S2A, S2B; SI**). Taken together, these results strongly suggest that the current generation of droplet-based scRNA-seq technologies (10x 3’ v3 and 5’) return far greater numbers of bacterial and viral UMIs than previous generation (10x 3’ v2).

#### Intracellular microbes can be distinguished from environmental contaminants in both plate and droplet-based scRNA-seq datasets

To rigorously distinguish contaminant taxa from true signal, we developed separate computational and statistical tests for plate-based (described in the **SI**) and droplet-based (10x 3’ v3 and 5’) scRNA-seq technologies. To control for (the low levels of) contamination in 10x 3’ v3 and 5’, we focus on genera with ≥ 2 reads per cell, which we derive from Robinson2023-10x where *Fusobacterium* was the only genus with at least 2 reads in a single cell across all conditions. We subsequently use the term “infected” to refer to any cell with ≥ 2 reads from the same genus because it strongly enriches for cells exposed to live bacteria (329/330 cells meeting this threshold in Robinson2023-10x were exposed to live *F. nucleatum*). We observed that infected cells are disproportionately either THP1 or HCT116 compared to Jurkat T from both the 3’ v3 (FDR=1.6e^-11^ (HCT116); FDR=3.1e^-13^ (THP1)) and 5’ datasets (FDR=0.0002 (HCT116); FDR=5.6e^-6^ (THP1)) (**Figure S2D, Table S1**). These approaches are also consistent for viruses, as we found *Hepatitis B* and *Merkel polyomavirus* to be the only infecting taxa (≥ 2 reads per cell) and found these viral taxa strongly enriched in tumor cells in Ma2021 (p-value=5e^-87^) and Paulson2018 (p-value=4e^-43^), datasets of hepatocellular and Merkel cell carcinoma, respectively (**Figures S2E, S2F, Table S1**).

In summary, intracellular microbes can be detected and distinguished from contaminants from both plate and droplet-based scRNA-seq technologies using cell type-specific and -agnostic approaches. For these reasons, we focus herewith on newer droplet-based scRNA-seq datasets (10x 3’ v3 and 5’) to interrogate the tumor microbiome of esophageal and colon cancer, which is the focus of the next part of the Results.

### B. Interrogating the intratumoral microbiome of colorectal and esophageal carcinomas Overview of the analysis

Having demonstrated that our pipeline correctly and specifically identified reads from *Fusobacterium* in a controlled experiment, we next sought to examine the intracellular tumor microbiome of patient tumors in publicly available datasets. We decided to focus on colorectal and esophageal cancers because there are available 10x 3’ v3 and 5’ scRNA-seq datasets with large numbers of patients, *Fusobacterium* has been previously reported to be highly abundant and be associated with worse prognosis in both cancer types, and the tumor microbiome of these cancer types has not been previously explored by scRNA-seq^9, 10, 34–37^. The colorectal carcinoma cohort (a subset of the above Pelka2021 dataset) includes 30 tumor samples and 4 adjacent non-tumor tissue samples from 20 tumors sequenced using 10x 3’ v3 (**Figure 3A**). The esophageal carcinoma cohort, Zhang2021, includes 110 tumor samples (divided into CD45-positive and negative cells) and 8 adjacent non-tumor tissue samples from 55 esophageal carcinomas sequenced using 10x 5’ (**Figure 3A**).

**Figure 3:**
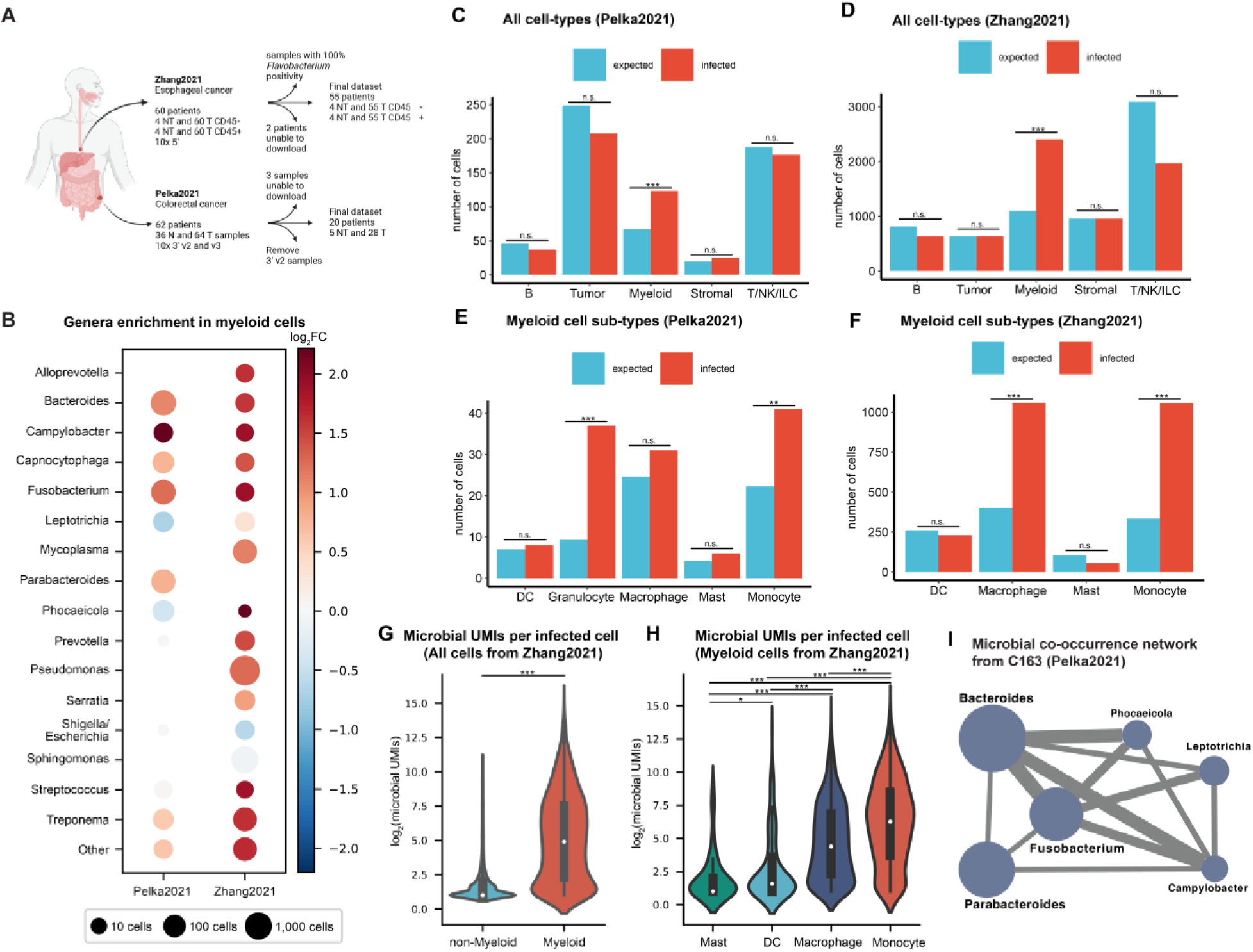
The cell type-specific bacterial landscape of CRC and ESCA patient tumors. **(A)** Overview of the samples analyzed in each of the publicly available scRNA-seq datasets. **(B)** The number of cells infected (≥ 2 UMIs) by each microbial genera where node size is relative to the number of infected cells (node sizes are normalized by the number of cells in each dataset) and node color indicates the log_2_FC in myeloid cells (> 0 indicates enrichment in myeloid cells) **(C)** The number of infected cells relative to expectation for the top-level cell types from Pelka2021. **(D)** The number of infected cells relative to expectation for the myeloid cell sub-types from Pelka2021. **(E)** The number of infected cells relative to expectation for the top-level cell types from Zhang2021. **(F)** The number of infected cells relative to expectation for the myeloid cell sub-types from Zhang2021. **(G)** The number of microbial UMIs identified per infected cell grouped by cell type (myeloid vs. non-myeloid) from Zhang2021. **(H)** The number of microbial UMIs identified per infected myeloid cell grouped by myeloid cell sub-type from Zhang2021. **(I)** The intracellular microbe co-occurrence network from colorectal tumor C163. Node size is relative to the number of cells infected by each microbe (Bacteroides is the largest node with n=160 infected cells). Edges appear between microbes that co-occur in the same cell more than expected (FDR < .05) and edge width is proportional to the statistical significance of the co-occurrence. * p-value < 0.05; ** p-value < 0.01; *** p-value < 0.001.

Next, we analyzed the tumor samples and observed a consistent range of the percentage of infected cells across patients in Pelka2021 (mean 0.7% and range 0-8.2% of the total number of cells) and Zhang2021 (mean 4.25% and range 0-21.7%), except for six tumor samples from three patients in Zhang2021 in which nearly 100% of cells were infected predominately by *Flavobacterium*, a previously reported cell culture contaminant^38^. After excluding these six contaminated samples, we found a total of 569 infected cells (0.7% of the cells sequenced) from Pelka2021 and 6,550 infected cells (3.4% of the cells sequenced) from Zhang2021 (see **Figure S3A, S3B** for UMAP of cell types and microbial UMIs). Fewer cells from the adjacent non-tumor tissue samples were infected compared to the matched tumor tissue samples in both datasets (0.1% of cells vs. 1.3% of cells from the paired non-tumor and tumor tissue respectively). Overall, we found 91 genera that infect at least one cell in either dataset (**Table S2**). Setting a threshold of ≥ 10 infected cells, we identified 16 commonly infecting genera, including eight genera found in both datasets, seven genera specific to Zhang2021 and one genus specific to Pelka2021 (**Figure 3B**).

#### Microbial UMIs are enriched in myeloid cells compared to other cell types in the TME

Given our previous *in vitro* finding that the proportion of bacterial read-positive cells can differ significantly between cell-lines (**Figure 1E**), we next asked whether specific cell types in the TME are also disproportionately infected. We found myeloid cells were the only cell type with significantly more infected cells than expected in both Pelka2021 (p-value=2e^-10^; log_2_FC=0.9) and Zhang2021 (p-value < 1e^-300^; log_2_FC=1.1) (**Figures 3C, 3D**; STAR **Methods**). Within the myeloid cell compartment, monocytes were the only myeloid cell sub-type infected significantly more than expected in both Pelka2021 (log_2_FC=0.88; p-value = 0.006) and Zhang2021 (log_2_FC=1.65; p-value = 5e^-247^) (**Figures 3E, 3F**). This enrichment in myeloid cells was not due to a small number of genera, as 13/16 of the commonly infecting genera and all other genera combined infect myeloid cells more than expected in at least one dataset (**Figure 3B**).

Next, we compared the number of microbial UMIs per infected cell across cell types and found that myeloid cells contained significantly more microbial UMIs compared to infected non-myeloid cells in both Pelka2021 (p-value = 0.048) and Zhang2021 (p-value < 1e^-300^) (**Figures 3G**, **S3C**). Within the myeloid cell compartment, infected monocyte cells contained significantly more microbial UMIs than the other myeloid cell sub-types in Zhang2021 (**Figure 3H**). These results showcase the importance of myeloid cells, particularly monocyte-derived cells, as the predominant source of bacterial presence in the intracellular tumor microbiome.

#### Bacterial genera exhibit specific co-occurrence relationships in the same cells

We observed that many more cells (n=64 and n=205 in Pelka2021 and Zhang2021, respectively) were infected by multiple bacterial genera than expected by chance alone (n=18 and n=108). This was true for both myeloid cells (n=10 and n=134 vs. n=3.5 and n=91 expected in Pelka2021 and Zhang2021, respectively) and non-myeloid cells (n=54 and n=64 vs. n=15 and n=34) in both datasets. As specific bacterial genera including *Fusobacterium*, *Leptotrichia* and *Campylobacter* have been described as co-enriched in colorectal tumors^39, 40^, we examined whether co-infection relationships between specific bacterial genera could be detected within single cells of individual tumors (STAR **Methods**). In total, we found 12 significant co-infection relationships in Pelka2021 and 9 significant co-infection relationships in Zhang2021 (FDR-corrected p-value < 0.05; **Table S3**). Occasionally, we observed infection of the same cell by three or more bacterial genera (n=19 and n=15 in Pelka2021 and Zhang2021). One extreme example of this phenomenon is colorectal tumor C163 where we found 14 cells co-infected by three or more genera, including a single tumor cell positive for six genera, including *Fusobacterium* (150 UMIs), *Campylobacter* (96 UMIs) and *Bacteroides* (42 UMIs) (**Figure 3I**). Our results suggest that bacterial genera may preferentially co-infect the same single cells in the tumor microenvironment, which agrees with a previous report that primary *F. nucleatum* infection greatly increased secondary invasion rates of other species *in vitro*^41^.

#### Intracellular bacteria induce the upregulation of antigen presentation genes in infected host tumor cells but their down-regulation in infected myeloid cells

Next, we sought to identify any host transcriptomic changes associated with the presence of intracellular bacteria in host cells. We first performed differential expression analysis between bacterial-positive and -negative cells for each cell type and each dataset separately. Intriguingly, we found that myeloid cells and tumor cells had many more differentially expressed genes (DEGs) between infected and bacterial-negative cells (mean 396 and 390 DEGs for myeloid and tumor cells respectively) compared to the other cell types (mean 68 DEGs) (**Table S4**). We also observed significantly more DEGs for *in vitro F. nucleatum*-exposed HCT116 epithelial cells (571 and 156 DEGs compared to unexposed cells and cells exposed to heat-killed *F. nucleatum,* respectively) and THP1 monocytic cells (1,985 and 249 DEGs) compared to Jurkat T cells (143 and 61 DEGs) from Robinson2023-10x (**Table S4**). Given these results, we focused our analysis on myeloid and tumor cells and performed gene set enrichment analysis (GSEA)^42^ on the DEGs identified between infected and bacterial-negative cells. We further clustered gene sets with overlapping genes using EnrichmentMap^43, 44^.

Interestingly, infected myeloid cells in both datasets shared a cluster of upregulated gene sets associated with response to molecules of bacterial origin, which points to the potential and consistent functional impact of these infections (**Figures 4A, 4B; Table S5**). In Pelka2021, we further observed one cluster of down-regulated gene sets associated with antigen processing and presentation, which has been previously reported as a mechanism of immune evasion by cells infected by bacteria^26, 45, 46^. We previously observed that monocytes are disproportionately infected relative to other myeloid cells in both datasets (**Figure 3E, 3F**), so we repeated our analysis comparing infected and bacteria-negative monocytes and found similar enriched pathways (**Figure S4A, S4B**), which suggests the identified enriched pathways are not due to different myeloid sub-type specificities.

**Figure 4:**
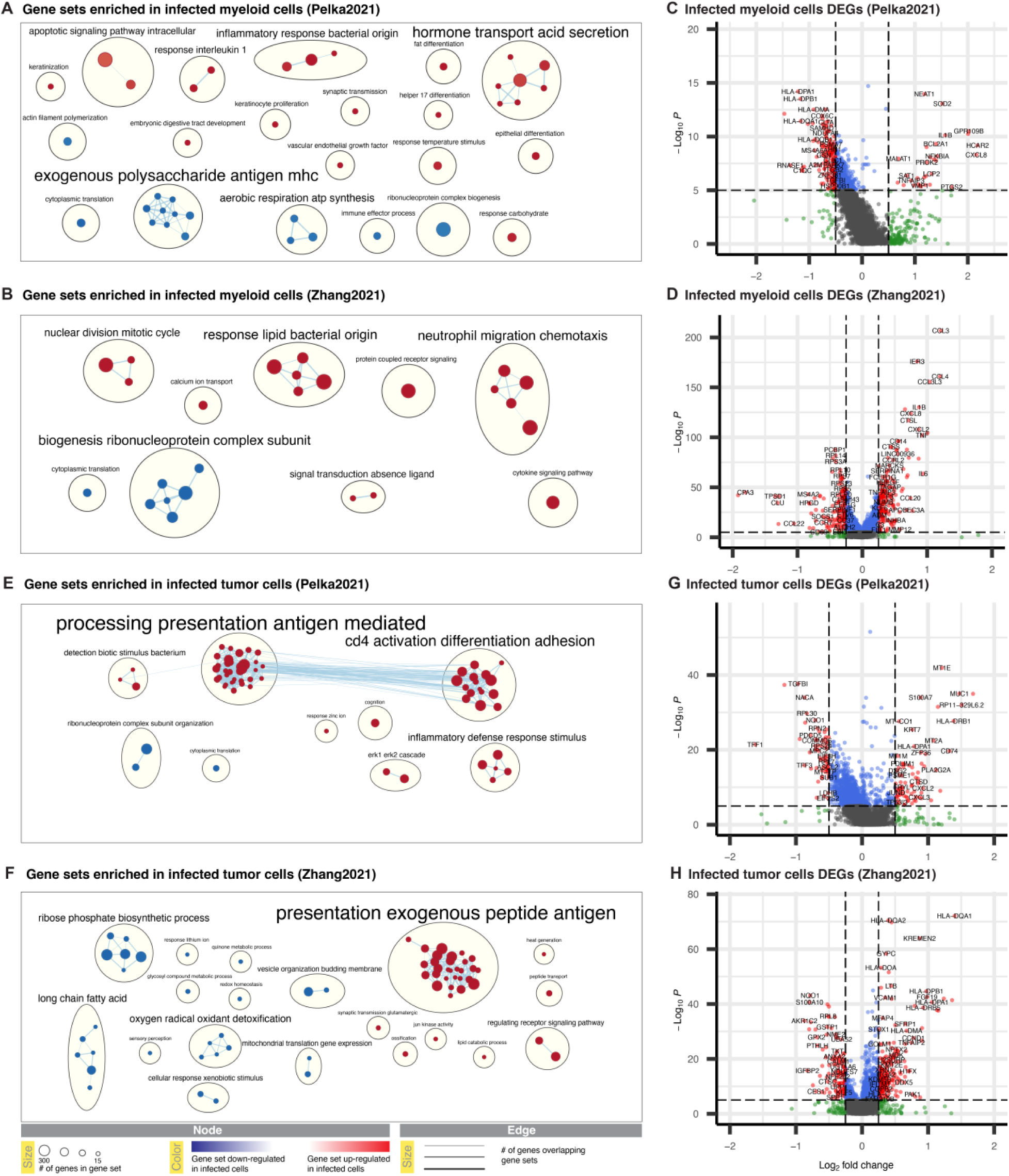
Host cell transcriptomic changes associated with bacterial infection. **(A)** EnrichmentMap of clustered gene ontology (GO) biological processes (BP) gene sets up and down-regulated in infected myeloid cells from Pelka2021. **(B)** EnrichmentMap of clustered GO BP gene sets up and down-regulated in infected myeloid cells from Zhang2021. **(C)** Volcano plot showing differentially expressed genes (DEGs) between “infected” and bacterial-negative myeloid cells from Pelka2021. **(D)** Volcano plot showing DEGs between “infected” and bacterial-negative myeloid cells from Zhang2021. **(E)** EnrichmentMap of clustered GO BP gene sets up and down-regulated in infected tumor cells from Pelka2021. **(B)** EnrichmentMap of clustered GO BP gene sets up and down-regulated in infected tumor cells from Zhang2021. **(C)** Volcano plot showing differentially expressed genes (DEGs) between “infected” and bacterial-negative tumor cells from Pelka2021. **(D)** Volcano plot showing DEGs between “infected” and bacterial-negative tumor cells from Zhang2021.

At the individual gene level, 15 genes were up-regulated in infected myeloid cells in both datasets (> 50-fold over-enrichment; p-value < 1e^-25^) and 33 were down-regulated in both datasets (> 8-fold over-enrichment; p-value < 1e^-20^) **(Figure 4C, 4D, S4C, S4D; Table S4**). These 15 shared up-regulated genes strongly overlapped with both the set of up-regulated genes in THP1 cells exposed to live *F. nucleatum* in Robinson2023-10x (12-fold over-enrichment; p-value < 4e^-13^) and the set of genes with NF-κB promoter sites (NF-κB target genes) (23-fold enrichment; p-value = 1e^-12^) (https://www.bu.edu/nf-kb/gene-resources/target-genes/) (**Figure S4C; Table S4**). NF-κB can be activated by several up-stream mechanisms including toll-like receptors (TLRs), which recognize conserved microbial molecules (PAMPs or pathogen-associated molecular patterns) including bacterial lipopolysaccharides (LPS)^47^. 15 of the top 16 abundant genera identified in this study are Gram-negative bacteria, which have LPS in their outer membranes. Of the 10 NF-κB target genes up-regulated in all datasets, *CXCL8* is potentially the most relevant clinically because high levels of interleukin 8 (IL-8) (the protein encoded by *CXCL8*) in the serum and the TME have been associated with a reduced clinical benefit from immune-checkpoint inhibitors across multiple cancer types^48, 49^. One of these previous reports found intratumoral IL-8 production was driven primarily by *CXCL8-*high myeloid cells that up-regulated IL-1 response genes (observed in both datasets) and down-regulated antigen processing and presentation (observed in Pelka2021) (**Figure 4A**) ^49^. Combining these findings suggests that bacterial infection of myeloid cells in the TME causes the production of IL-8, which decreases response to immune checkpoint blockade, a hypothesis for future experimental investigation.

Beyond the binary differential expression analysis, we further hypothesized that the actual number of microbial UMIs would be a direct function of the number of infecting microbes and these numbers may be associated with host cell gene expression changes. We identified genes whose expression significantly correlated with the number of microbial UMIs in myeloid cells (excluding bacteria-negative myeloid cells) in both datasets (**Table S4**). Reassuringly, we found the correlation rho values for dysregulated genes between Pelka2021 and Zhang2021 (Spearman Rank Correlation rho=0.14, p-value < 2e^-58^) and observed that previously discussed genes were either positively correlated (*CXCL8* and *IL1B*) or negatively correlated (many HLA-II genes) with microbial load in both datasets. We then repeated this analysis using only one genus (*Pseudomonas*, which is the most abundant genus in Zhang2021) and identified similar results (**Table S4**). These results emphasize “microbial load” as an important factor in modulating host cell gene expression, something to be considered in future scRNA-seq-based explorations of the tumor microbiome.

Next, we analyzed tumor cells and observed the largest cluster of up-regulated gene sets in infected tumor cells to be associated with antigen processing and presentation and cell killing by leukocytes in both Pelka2021 and Zhang2021 (**Figure 4E, 4F**). At the individual gene level, we found 23 genes up-regulated in infected tumor cells in both cancer-types (13-fold over-enrichment; p-value < 1e^-19^), including the cytoskeletal gene *KRT7*, which is up-regulated by *F. nucleatum* infection and promotes metastasis of colorectal carcinoma^50^, and 12 genes involved in antigen processing and presentation of peptides on HLA-I and -II (see **Figure S4E, S4F** for complete list). The up regulation of HLA genes by infected tumor cells is surprising on two levels. First, intracellular bacteria have been reported to decrease HLA expression levels in host cells to evade the immune system^26^, which we do observe specifically in the myeloid cells. Second, tumor cells also have frequent HLA alterations including HLA loss-of-heterozygosity (LOH)^51, 52^ and HLA down-regulation^53^ to evade the immune system. While the idea that tumor cells and intracellular bacteria may cooperate to evade the immune system together is intuitive, our transcriptomic data suggests that tumor cells respond to bacterial invasion by up-regulating antigen processing and presentation, which may increase the presentation of neo-antigens and lead to tumor killing by the immune system.

Collectively, our analyses of the proportion of infected cells, the number of microbial UMIs per infected cell, and host transcriptomic changes associated with infection from patient tumor scRNA-seq datasets, suggests a model of the tumor microbiome. In this model, bacteria predominately reside within co-opted myeloid cells that trigger inflammatory pathways like TNF and IL-1 via NF-κB pathway activation while simultaneously shielding the bacteria in these cells by down-regulating antigen processing and presentation pathways. In contrast, while bacterial UMIs are found in tumor cells and strongly enriched in tumor samples (compared to non-tumor adjacent tissue), they do not appear to preferentially infect tumor cells by either the proportion of infected cells or the number of microbial UMIs per infected cell. One potential explanation for this is the upregulation of HLA genes in infected tumor cells, which suggests that tumor cells respond to bacterial invasion by “raising the alarm” even at risk of being detected and killed by the immune system.

## Discussion

In this study, we sought to identify microbial reads via the analysis of scRNA-seq data. Our first main goal was a methodological one, to systematically study the detection capabilities of such an approach across an array of different sequencing platforms. Due to the lack of suitable published positive control datasets, we generated a new dataset of cell lines of distinct lineages exposed to live and heat-killed bacteria using both plate and droplet-based scRNA-seq approaches. By analyzing this dataset (Robinson2023) along with other related publicly available datasets, we systematically identified important differences in the number of bacterial reads sequenced from known intracellular microbes and contaminants, both between plate and droplet-based approaches and further, within droplet-based approaches (10x 3’ v2 vs. 3’ v3 and 5’). The increased number of contaminating reads observed from plate-based scRNA-seq datasets is not surprising as it is likely easier for contaminants to enter wells compared to microfluid droplets.

The difference in microbial reads across 10x chemistries sheds further light on the distinct approaches to contamination taken by alternative existing scRNA-seq microbiome pipelines like SAHMI and INVADEseq. SAHMI analysis of 10x 3’ v2 pancreatic datasets would have necessitated multiple decontamination protocols^19^, given the relatively poor microbial read recovery of this assay. While the INVADEseq study does not discuss contamination, it sets a comparable threshold (≥ 3 reads vs. ≥ 2 reads, respectively) for host cell differential expression analysis for their 10x 5’ assay custom modified to detect bacterial 16S rRNA transcripts^21^. Importantly, our analysis suggests an intermediate stringency with which unmodified 10x 3’ v3 and 5’ datasets (many of which have already been generated) can be analyzed for cell host-associated microbes using a simple threshold for removing contaminants.

Based on the findings related to our first, methodological analysis, we turned to our second, thematic goal: analyzing the intracellular microbiome of esophageal and colorectal carcinoma, to answer two basic questions: (a) what is the cell type-specific tumor microbiome landscape in these tumors? and (b) what are the cell type-specific host transcriptomic changes that are associated with these infections? In response to the first question, we find that intracellular microbes can be found predominately (but not exclusively) within myeloid cells in the TME and that some intracellular bacteria co-occur within the same cells in the TME. Answering the second question, we find that infected myeloid and tumor cells have cell type-specific transcriptomic alterations with potential functional impact.

Our analysis of the host transcriptomic changes in infected cells suggests that bacteria may play an important role in modulating immune processes, which suggests intratumoral microbial load may impact efficacy of cancer immunotherapy. Infected myeloid cells appear to significantly up-regulate expression of *CXCL8* (and likely production of IL-8), and high levels of IL-8 have been reported to be associated with decreased response to immune checkpoint blockade^49^. In contrast to infected myeloid cells, infected tumor cells transcriptionally up-regulate antigen processing and presentation and other pathways that may make them more susceptible to killing by T cells. These results suggest that not only microbial load, but also type of infected cell, are important considering factors in designing immunotherapy regimens.

Our results suggest several hypothetical clinical approaches to modulate intratumoral microbes for improved therapeutic results. One approach would be to introduce invasive bacteria such as *Salmonella* to prime tumor cells for killing before immunotherapy, as previously described^54, 55^, but this may be most beneficial in tumors with low myeloid TME infiltration. An inverse approach would be to target intracellular bacteria in myeloid-high tumors, by reducing infected myeloid cells and subsequently IL-8 levels, which should lead to improved response to immunotherapy. However, targeting intracellular bacteria is difficult because many antibiotics do not penetrate the membrane of host cells^56^. Even chemotherapeutic drugs like 5-fluorouracil, which penetrates host cell membranes and inhibits *Fusobacterium* growth, can be modified to a non-toxic form by other members of the intratumor microbiota including *Bacteroides*^57^, which we find to be the most common co-infecting genera with *Fusobacterium* in Pelka2021. Given these complications, it may be preferable to target the downstream signaling pathways associated with infected myeloid cells, e.g., by using an IL-8 or NF-κB inhibitor^58, 59^.

There are several important limitations of our study, most notably our hypothesis that the microbial UMIs we identified across scRNA-seq datasets are indicative of viable intracellular bacteria. First, our arbitrary cut-off of ≥2 UMIs may not be sufficient to distinguish “infected” from “bystander” cells, which is supported by the correlation of the expression of genes with microbial load in myeloid cells in both datasets. Second, reads may emanate from phagocytized (and killed) bacteria, particularly in myeloid cells. However, our *in vitro* model suggests this is unlikely to occur at high rates, as we observed only one cell with ≥2 *Fusobacterium* UMIs that was exposed to heat-killed *F. nucleatum*; all other cells were exposed to live bacteria. Third, one cannot rule out that at least some of the polymicrobial communities we describe as intracellular are actually extracellular but adhere to the cell surface via biofilm formation. Indeed, polymicrobial biofilms featuring *Fusobacterium* and *Bacteroides* have been described in colorectal cancer^60^, but we note that if indeed this sometimes occurs, the biofilms interestingly are created in a cell-type-specific manner. Fourth, microbial contaminants may be introduced during laboratory manipulation of tissue samples, which we find evidence of in the esophageal dataset. Indeed, nearly 100% of cells showed infection in three patients and the identified genus was common between all three samples. Although this is to the best of our knowledge the first report of contamination within scRNA-seq data, we note that laboratory contamination has been recognized as a significant issue in cell cultures that can have important implications for cell phenotype and biology^11^.

In summary, we have performed the first systematic evaluation of multiple scRNA-seq technologies for their efficacy in recovering microbial UMIs. Based on the findings of the latter, we analyzed two pertaining publicly available patient datasets to chart their tumor microbiome. The latter analysis has identified co-abundant microbes associated within individual myeloid and tumor cells. Our analysis points to two potentially opposing effects of microbial infection on the immune susceptibility of tumors: while intracellular bacteria residing within myeloid cells may decrease their immune susceptibility, intracellular bacteria residing within tumor cells may increase the latter. Notably, the expression alterations observed in tumor cells are remarkably consistent across *in vitro* and patient tumor datasets. Overall, these findings suggest that by manipulating the intratumoral microbiome, for example via targeted antibiotics, one may hope to improve response to immune checkpoint blockade treatments, but that the overall functional effects of such future treatments are likely to depend on the level of myeloid infiltration in the TME in a complex manner.

## Acknowledgements

Support from CCR Single Cell Analysis Facility for single cell RNA-Seq captures, library preparation, sequencing and primary data processing was funded by FNLCR Contract 75N91019D00024. This work utilized the computational resources of the NIH HPC Biowulf cluster. (http://hpc.nih.gov) This research was supported in part by the Intramural Research Program of the National Institutes of Health, National Cancer Institute. WR’s contribution to this research was supported in part by NSF award DGE-1632976. The authors thank Wolfgang Resch for assistance running this analysis on the NIH HPC Biowulf cluster. The authors thank Dr. Ken-ichi Hanada for the gift of the Jurkat T cell line. The authors thank Anna Aulicino, Noa Bossel Ben-Moshe, Leigh Greathouse, Hector Corrado-Bravo, Norma Andrews, Steve Christensen, Itay Tirosh, Livnat Jerby-Arnon, Yunhua Zhu, Chen Zhao, Spyros Darmanis, Frank Lowery and Sri Krishna for useful discussions about this project.

## STAR Methods

### Resource Availability

#### Lead contact

Further information and requests for resources and reagents should be directed to and will be fulfilled by the lead contact, Eytan Ruppin.

### Materials availability

This study did not generate new unique reagents.

### Data and code availability

Single-cell RNA-seq data generated by this study is in the process of being deposited at GEO and will be publicly available as of the date of publication. Accession numbers will be listed in the key resources table.

Our pipeline is made publicly available on GitHub and uses Snakemake^61^ and conda, which are workflow and package management systems, to enable: reproducibility of our results, integration with high performance computing systems and analysis of microbial reads from additional scRNA-seq datasets by other researchers. CSI-Microbes is a bioinformatics pipeline combining several existing methods for sequence analysis with new methods for statistical analysis. CSI-Microbes includes considerable software engineering and use of Snakemake to connect the methods robustly and to enable the connected methods to run efficiently in different computing environments.

All original code has been deposited at Zenodo and is publicly available as of the date of publication. DOIs are listed in the key resources table. CSI-Microbes is logically partitioned into two modules, one module for the “identification” step and one module for the “analysis” and “validation” steps. A reproducible Snakemake^61^ (https://snakemake.readthedocs.io/) workflow for identifying microbial reads from scRNA-seq datasets, which includes code to download the data from the datasets above, is available on GitHub (https://github.com/ruppinlab/CSI-Microbes-identification) although the identification module has some dependencies to the NIH Biowulf server. A reproducible Snakemake workflow for analyzing microbial reads to identify intracellular microbes is available on GitHub (https://github.com/ruppinlab/CSI-Microbes-analysis). To facilitate reproduction of our analyses, we have uploaded CSI-Microbes-analysis v1.0.0 (the version used in this study) along with the relevant data files (generated by CSI-Microbes-identification v1.0.0) to Zenodo (https://doi.org/10.5281/zenodo.4695248).

### Experimental Model and Subject Details

#### Cell and bacterial culture

HCT116 (colorectal carcinoma) cells were grown in McCoy’s 5A Media containing 10% FBS (Corning, #10-050-CV). Jurkat T (T cell acute lymphoblastic leukemia) and THP-1 (acute monocytic leukemia) cells were each grown in RPMI-1640 containing 10% FBS (Corning, #10-040-CV). All cell lines were maintained in a 37°C degree incubator with 5% CO2. *Fusobacterium nucleatum* ATCC 25586 was cultured anaerobically in Columbia broth (BD Difco, #294420) at 37°C with 200 rpm shaking. All cell lines and bacterial strains were obtained from the ATCC (Manassas, VA). Cell line identities were authenticated by the ATCC and cells screened negative for *Mycoplasma* contamination.

### Method Details

#### Invasion assay

Cell lines were seeded at a density of 10^6^ prior to infection. Bacteria were grown to late log phase and then 5 µM CellTrace Violet (ThermoFisher, #C34571) was added to the broth culture for 20 minutes with shaking. Bacteria were pelleted and washed in sterile 1X PBS (Corning, #21-031-CV), then added to cell culture media at an MOI of 100:1. Cells were infected for 10 hours then media was removed and cells were washed twice with 1X PBS. Fresh media containing 200 µg/mL gentamicin (ThermoFisher, #15750060) and 200 µg/mL metronidazole (ThermoFisher, #AC210340050) for 1-2 hours to kill extracellular bacteria. After antibody incubation, cells were washed twice with sterile X PBS then collected for FACS analysis by trypsinization (HCT116 only). Bacterial invasion efficiency was determined by serial dilution and plating on TSA + 5% sheep’s blood (Remel, #R01200) following a 10-minute incubation of cells in 1% Triton X-100 (Sigma Aldrich, #T8787). Agar plates were incubated anaerobically for 72 hours prior to colony enumeration.

#### Droplet-Based Single Cell Partitioning, Library Preparation and Sequencing (10X Genomics)

Single cell suspensions were prepared, and concentration and viability measured on an automated dual-fluorescence cell counter with acridine orange and propidium iodide stain (Luna Fx7, Logos Biosystems). Single cell partitioning and RNA-Seq library preparation was performed using 10X Genomics Chromium NextGEM 5’ v2 chemistry (user guide CG000331), or 10X genomics 3’ v3.1 chemistry (user guide CG000204) according to vendor recommendations. Sample viability was above 80% for all conditions. 6,000-8,000 cells were targeted to be captured for each sample. Libraries were sequenced on the NovaSeq 6000 with a target depth of 50,000 reads per cell using read parameters recommended by 10x Genomics user guides. Reads from multiplexed sequencing runs were demultiplexed using *cellranger mkfastq* v7.0.0 (10x Genomics).

#### Plate-Based Single Cell Partitioning, Library Preparation and Sequencing (SeqWell PlexWell)

Single cell suspensions were prepared and sorted on a BD FACS Aria IIU into 96 well PCR plates that were prepared with lysis buffer according to seqWell plexWell Rapid Single Cell method (user guide v20210402) and containing ERCC spike-in mix (Invitrogen # 4456740) at a dilution of 1:1E7. Following deposition of single cells into prepared plates, they were snap-frozen and stored at -80C until further processing. Single cell cDNA and libraries were generated for each sorted sample according to seqWell plexWell Rapid Single Cell user guide. Multiplexed libraries containing indexed single cell RNA-Seq libraries were sequenced on the either the NextSeq 2000 or NovaSeq 6000 with a target read depth of 2 million reads per cell using the following read parameters: Read 1 61bp, Index 1 8bp, Index 2 8bp, Read 2 61bp. Raw sequencing data was demultiplexed into individual sample fastq sets using *bcl2fastq v2.20.0*.

#### Plate-Based Single Cell Dataset Alignment to Human Genome and Analysis

Raw FASTQ files were trimmed using fastp^62^ v0.20.1 with the arguments “*--unqualified_percent_limit 40 --cut_tail --low_complexity_filter --trim_poly_x*”. The trimmed FASTQ files were aligned to the reference human genome (GRCh38 gencode release 34) and any applicable spike-in sequences using STAR^63^ 2.7.6a_patch_2020-11-16 with the arguments “--soloType SmartSeq --soloUMIdedup Exact --soloStrand Unstranded --outSAMunmapped Within”. For analysis, we used the previously described workflow “Lun 416B cell line (Smart-seq2)” from “Orchestrating Single-Cell Analysis with Bioconductor” (http://bioconductor.org/books/3.16/OSCA.workflows/lun-416b-cell-line-smart-seq2.html#lun-416b-cell-line-smart-seq2) to specifically handle ERCC spike-in sequences. Quality filtering was performed using the quickPerCellQC function from scuttle (citation; version 1.8.0).

#### 10x Single Cell Dataset Alignment to Human Genome and Analysis

Raw FASTQ files were aligned to the reference human genome (GRCh38 gencode release 34) using CellRanger^64^ v5.0.1 (https://support.10xgenomics.com/single-cell-gene-expression/software/pipelines/latest/what-is-cell-ranger). The annotated polyA and template sequence oligonucleotide (TSO) sequences were trimmed, the unmapped reads were converted to the FASTQ file format trimmed and filtered using FASTP as described above before being converted to BAM files. For quality filtering, we removed genes found in fewer than 3 cells and remove cells with fewer than 3,000 gene features or greater than 15% of reads mapping to mitochondrial genes.

#### Annotation of cell types

For the Plex-Well dataset generated by this study, we sorted cells with known identity (HCT116 or Jurkat T) into wells or intentionally left wells empty. For the 10x datasets, we performed low resolution clustering using Seurat (resolution=0.1) and annotated clusters using marker genes for HCT116 (*EPCAM*), Jurkat T (*CD3E*) and THP1 (*CD64*/*FCGR1A*).

For previously generated datasets, we used the cell type annotations provided by the original authors. We harmonized annotations between the esophageal (Zhang2021) and colorectal (Pelka2021) datasets when they differed. From the esophageal dataset, we grouped endothelial, FRC, fibroblast and pericytes as Stromal cell (the top-level annotation for these cell types used in the colorectal carcinoma dataset). From the colorectal dataset, we grouped plasma cells as B cells and mast cells as myeloid cells using the author-supplied the top-level annotations. The cell type annotation harmonization resulted in five major cell type groupings: Epithelial (Tumor), Stromal, B, T/NK/ILC and Myeloid.

#### Alignment of unmapped reads to microbial genomes

The unaligned reads were assigned to microbial taxa using PathSeq^23^ v4.1.8.1 (http://software.broadinstitute.org/pathseq/) with the arguments “*--filter-duplicates false --min-score-identity .7*”. We constructed the reference microbial genome database by downloading the set of complete viral, bacterial and fungal genomes from RefSeq release 201^65^. We subsampled at least one genome from each species including any genomes annotated as either “reference genome” or “representative genome” as well as the genomes of the three *Salmonella* strains used in the analyzed datasets. To mitigate vector contamination, we identified regions of suspected vector contamination (including “weak” matches) in the genomes using Vecscreen_plus_taxonomy (https://github.com/aaschaffer/vecscreen_plus_taxonomy) with the UniVec Database (ftp://ftp.ncbi.nlm.nih.gov/pub/UniVec/) and filtered any reads that aligned to these regions^66^.

### Quantification and Statistical Analysis

#### Single sample cell type enrichment (plate-based protocols)

We define the abundance of a particular microbe in each cell to be the number of unambiguous reads assigned to the relevant genome(s) by PathSeq The abundances are normalized using the computeSpikeFactors function from scran^67^ v1.16.0 (https://github.com/MarioniLab/scran), which computes the library size factors using the sum of the spike-in sequences. To limit the number of hypotheses, we only test microbial taxa with counts per million microbial reads > 10 in at least 50% of the cells from a cell type. The logged normalized read counts are compared across cell types using the findMarkers function from scran v1.16.0 with arguments “*test=’wilcox’, lfc=0.5, block=’plate’*”. The findMarkers function from scran v1.16.0 makes inconsistent assumptions about how to distribute the values of the null distribution depending on whether the user specifies *“direction = ‘up’”* or *“direction = ‘down’”* (a one-sided test) or the user specifies *“direction = ‘any’”* when the parameter *lfc* is greater than zero (https://github.com/MarioniLab/scran/issues/86). The assumption for the one-sided test models our intent so we ran the comparison twice, once using with “*direction=’up’”* and once with “*direction=’down’”*, selected the result with the smaller p-value for each microbial taxa and converted the one-sided p-value to the two-sided p-value by taking the minimum of 1 and 2*p-value as described in a standard reference^68^ (page 79).

#### Single sample cell type enrichment (10x protocols)

We define a cell to be “infected” by a particular microbial taxon if there are ≥ 2 UMIs assigned unambiguously to the relevant genome(s) by PathSeq. We analyze only the cell type enrichment of microbial taxa that infect at least five cells in a given sample. We compare the percentage of cells infected by a given microbial taxa between two cell types in a given sample using the fisher_test function with argument ‘alternative=”greater”’ from SciPy).

#### Calculation of Number of Expected Infected Cells per Cell Type (10x protocols)

We first calculate the number of expected infected cells for a given cell type for an individual sample using the formula:

(number of cells from that cell type/number of cells) * number of infected cells

Next, we summed the number of cells expected to be infected for a given cell type across all samples and compared it to the actual number of infected cells for a given cell type. We report the value: log2(number of infected cells per cell type/ expected number of infected cells per cell type)

To calculate the significance of the difference between the number of expected and actual infected cell per cell type, we first calculate the per sample p-value for each cell type (see above). Next, for a given cell type, we combine the per sample p-values using Stouffer’s Z-score method (using the combine_pvalues function with argument ‘method=”stouffer”’ from SciPy) weighting the p-values using the number of expected infected cells (as samples with very few infected cells or very few cells of the cell type of interest will not be informative). We set the maximum per sample p-value to be .9999 as combine_pvalues function with method=”stouffer” returns (-Inf, 1) when any p-value = 1 (https://github.com/scipy/scipy/issues/8506).

#### Calculation of co-infection relationships

In each dataset (Pelka2021 and Zhang2021), we calculated co-infection relationships only for genera that infect ≥ 10 cells. We initially calculate co-infection relationships per sample. Given genera g1 and genera g2, we calculate the co-infection relationship using the hypergeometric enrichment to ask whether the number of cells infected by both g1 and g2 are more than expected knowing the total number of cells, the number of cells infected by g1 and the number of cells infected by g2 (using the hypergeom function from SciPy). Next, for a given co-infection relationship, we combine the per sample p-values using Stouffer’s Z-score method (using the combine_pvalues function with argument ‘method=”stouffer”’ from SciPy) weighting the p-values using the number of expected co-infected cells (as samples with very few infected cells of g1 or g2 will not be informative). We set the maximum per sample p-value to be .9999 as combine_pvalues function with method=”stouffer” returns (-Inf, 1) when any p-value = 1 (https://github.com/scipy/scipy/issues/8506).

#### Host cell transcriptomic analysis

We performed differential gene expression analysis between “infected” (≥2 microbial UMIs) and bacterial-negative (0 microbial UMIs) human cells for myeloid and tumor cells. We used the function FindMarkers from Seurat (v4.3.0) using the default value for the host transcriptomic results.

#### Gene set enrichment analysis and visualization

For gene set enrichment analysis (GSEA), we performed differential gene expression analysis as described above using the additional argument of “logfc.threshold = -Inf, min.pct = -Inf, min.diff.pct = -Inf” to return a p-value and avg_log2FC for every gene. We ranked genes using the formula: avg_log_2_FC*-log_10_(p-value + 1e^-300^). We add 1e^-300^ to avoid errors when the p-value equaled to zero. We ran GSEAPreranked (v4.3.2) using pre-ranked genes (described above) and the Gene Ontology Biological Processes v.7.5.1 gene sets using the default parameters and seed = 0.

We visualized the enriched gene sets using Cytoscape^69^ (v3.9.1) and Enrichment Map^44^ (v3.3.5) (https://enrichmentmap.readthedocs.io/) with parameters: p-value < .001 (node cutoff) and Jacard Overlap Combined Index (k constant=0.5) > .5 (edge cutoff). Clustering was performed on the graph using MCL Cluster from AutoAnnotate^70, 71^ (v1.4.0). Annotations were manually reviewed and edited where appropriate.

#### Correlation of host gene expression and microbial load

We used the R function corr.test (with the argument method=”spearman”) to identify correlations between normalized expression for all genes and the number of genera-resolution microbial UMIs. We excluded all cells with zero microbial UMIs to focus on microbial load and avoid identifying genes simply associated with infection. We used the R function corr.test (with the argument method=”spearman”) to compare the correlations for overlapping genes in Pelka2021 and Zhang2021.

