## Supplementary Information for "scRNA-seq analysis of colon and esophageal tumors uncovers abundant microbial reads in myeloid cells undergoing proinflammatory transcriptional alterations"

### Analysis of bacterial read type from different scRNA-seq technologies

One open question in the field of scRNA-seq microbiome studies is how bacterial reads are captured by polyA approaches (like 10x and plexWell) because bacterial reads are poly-adenylated at a much lower frequency and with significantly shorter polyA tails compared to eukaryotic mRNA[30]. To identify the type of bacterial transcripts captured by scRNA-seq technologies and whether this differs between technologies, we reanalyzed Robinson2023 using CSI-Microbes, using SRPRISM with default settings as the alignment tool against *F. nucleatum* subspecies *polymorphum* (Strain: NCTC10562). This was done because SRPRISM is designed to align sequences to a reference genome with a deterministic, user-controllable parameterization of how many nucleotide mismatches are allowed. We found that 82%, 92% and 95% of plexWell, 10x 5' v2 and 10x 3' v3 *F. nucleatum* reads mapped to rRNA regions (**Figure S1C**).

### Analysis of experimental carryover from different scRNA-seq technologies

We sought to compare the potential for experimental carryover, by examining the number of reads mapping to *Fusobacterium* in unexposed cells and empty wells. We examined the number of reads mapping to *Fusobacterium* and found zero reads in the unexposed control cells sequenced using 10x compared to 292 reads in the unexposed control cells and empty wells sequenced using plate-based technologies (**Figures 1D, 1E**). We confirmed this pattern in the *Salmonella* datasets (**Figures 1A, 1B**). We hypothesize that these *Fusobacterium* reads represent cross-well contamination as we sequenced unexposed cells and cells exposed to *F. nucleatum* in the same plate. We then examined the number of *Fusobacterium* reads from cells exposed to heat-killed bacteria and found significantly fewer compared to cells exposed to live bacteria

(**Figures 1D, 1E**). Taken together, these results testify that microbial UMIs captured by scRNA-seq technologies (particularly by droplet-based technologies) are mostly specific to live invasive bacteria, in agreement with a previous report[21].

### Analysis of viral reads from different 10x chemistries

In this study, we analyzed multiple scRNA-seq data sets (**Table 1**) generated using different methods, including three different 10x protocols: 3’ v2, 3’ v3, and 5’. To evaluate whether the quantities of viral reads detected differ substantially among these protocols, we compared data generated using more than one 10x protocol after processing with our CSI-Microbes pipeline. One limitation of this analysis is that we analyzed either samples at different timepoints from the same tumor (Ma2019) or samples from the same timepoint from different tumors (Paulson2018). We first analyzed Paulson2018, which is a 10x 3’ v2 and 5’ dataset of two Merkel cell carcinomas, caused by Merkel polyomavirus integration[33]. Even though the Merkel cell polyomavirus must undergo a substantial mutation before integrating into the host genome to cause cancer[34], we identified 11,141 Merkel polyomavirus reads from the tumor sequenced using 10x 5’ but only 691 and 394 reads from the two tumor samples sequenced using 10x 3’ v2 (p-value < $2.2e^{-16}$; odds ratio=3.13 and 4.16 respectively after correcting for sequencing depth) (Figure 1C). We next analyzed Ma2019, which is an scRNA-seq liver cancer dataset predominately sequenced using 10x 3’ v2 that happens to include one hepatocellular carcinoma with different timepoints sequenced using either 10x 3’ v2 or v3 with detectable reads from Hepatitis B in both samples [31,32]. Like our previous findings, we observed significantly more reads to Hepatitis B in the sample sequenced using 10x 3’ v3 compared to 3’ v2 in both absolute number (683 vs. 56 reads) and after controlling for sequencing depth (p-value < $2.2e^{-16}$; odds ratio=5.14) (Figure 1C). Our results, when taken together, strongly suggest that the current generation of droplet-based scRNA-seq technologies (10x 3' v3 and 5') return far greater numbers of viral UMIs than previous generation (10x 3' v2).

### Statistical Tests for Contaminants in Plate-based scRNA-seq Technologies

In Aulicino2018, we observed that *Salmonella* is the only genus that is significantly higher (FDR-corrected p-value < .05 with minimum $\log_2$FC = 0.5) in infected cells compared to bystander or unexposed control cells in Aulicino2018 despite the high number of reads detected from other microbial genera (FDR=1.4e$^{-7}$, **Figure 1A**, **Methods**, **Table S1**). We prospectively validated this approach on Robinson2023-plexWell, which only identified *Fusobacterium* both when comparing between different cell types (HCT116 and Jurkat) exposed to live *F. nucleatum* (FDR=2e$^{-8}$, **Figure 1D**, **Table S1**) and when comparing all cells exposed to live *F. nucleatum* to empty wells (FDR=0.013, **Figure S2C**, **Table S1**).

We note there are three important limitations of identifying intracellular microbes from plate-based datasets. The first limitation is that our cell type-agnostic approach for plate-based scRNA-seq technologies uses empty wells, which are important negative controls for microbiome studies. Unfortunately, those are not included in many plate-based scRNA-seq cancer datasets. The second limitation is the identification of many reads from the target microbe in cells without live bacteria (some of which likely result from cross-well contamination), which makes it less suitable than droplet-based technologies to a simple threshold approach. The third limitation is the comparison of the distribution of reads within cells, which will work best when most cells have reads from one genus and may miss genera that infect a small number of cells.

### Application of statistical tests to viral-infected cancers

We applied our two statistical tests for analyzing intracellular microbes from 10x data to the previously discussed 10x 3' v3 and 5' samples from Paulson2018 and Ma2019. The UMI-based filter ($\geq 2$ UMIs from the same genera per cell) identified *Merkel polyomavirus* and *Hepatitis B* as the only viral genera to infect any cells in Paulson2018 and Ma2019 respectively. Our cell type-specificity test found that infected cells were disproportionately tumor cells (Merkel polyomavirus FDR=$4.2e^{-43}$; Hepatitis B virus FDR=$5.4e^{-87}$), which is expected as both viruses have been shown to integrate into tumor cells (**Figure S2E, S2F**).
