## Supplementary Figures for "scRNA-seq analysis of colon and esophageal tumors uncovers abundant microbial reads in myeloid cells undergoing proinflammatory transcriptional alterations"

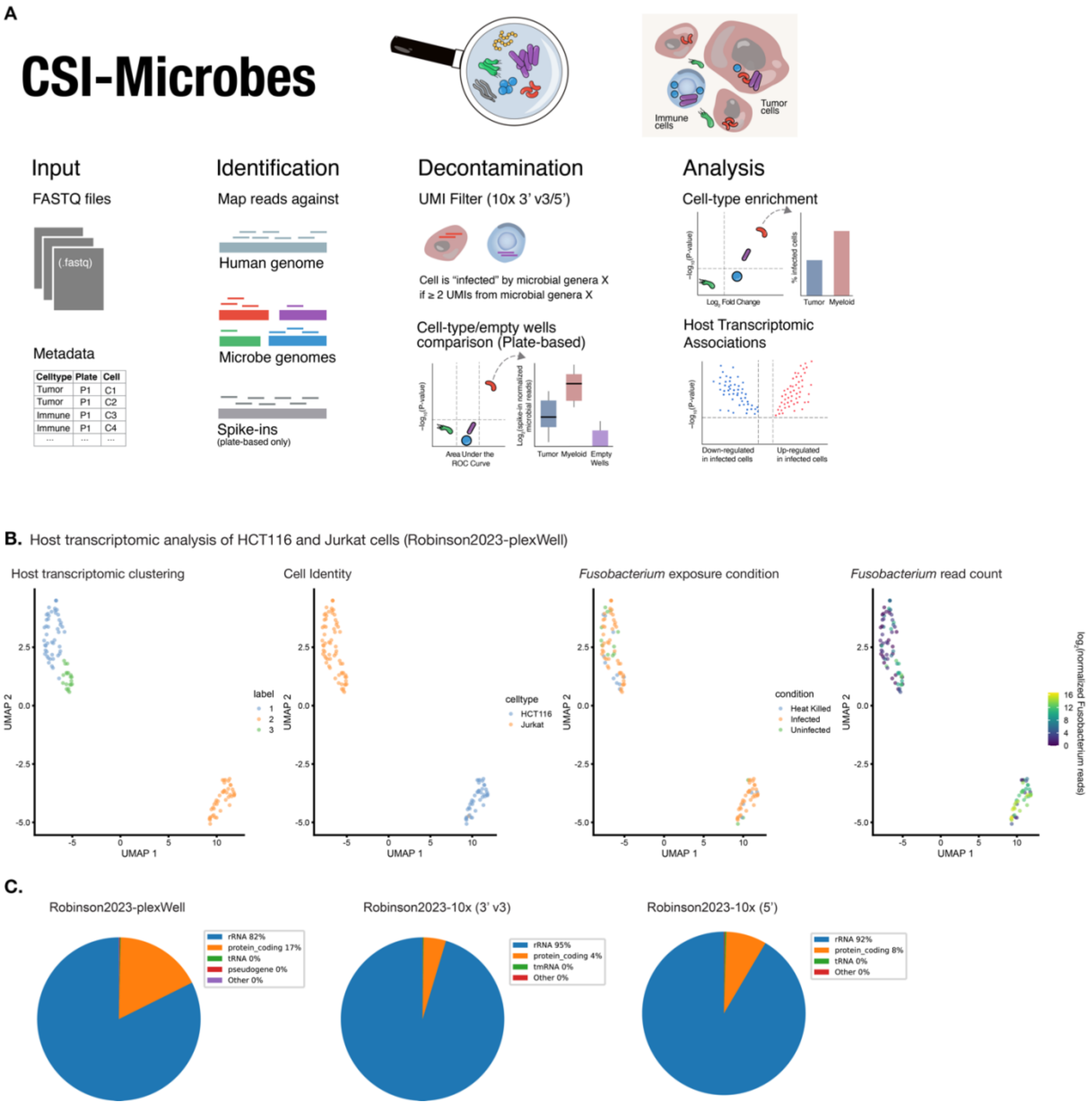


***Figure S1: A deeper analysis of CSI-Microbes and Fusobacterium infection and reads from Robinson2023***

***(A)*** *Overview of the steps in the CSI-Microbes pipeline including input, alignment, decontamination and analysis.* ***(B)*** *UMAP of HCT116 and Jurkat T cells sequenced using plate-based approaches colored by host transcriptomic clustering, cell identity, F. nucleatum exposure condition and the number of Fusobacterium reads identified per cell.* ***(C)*** *The proportion of Fusobacterium-aligned reads that map to different classes of RNA across the three different scRNA-seq protocols used in Robinson2023.*

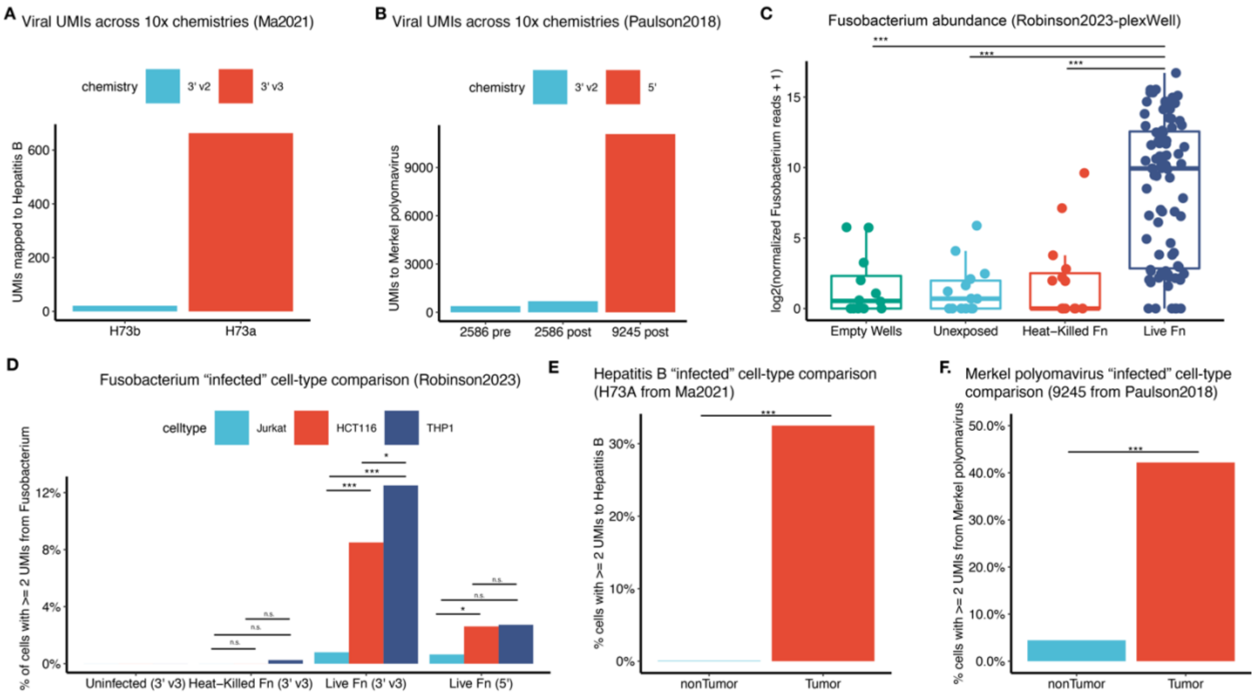


***Figure S2: CSI-Microbes identifies viral read detection differences and known microbial cell-type enrichments***

***(A)*** *The number of UMIs mapped to Hepatitis B from two timepoints from the same hepatocellular carcinoma sequenced using 10x 3' v2 and 10x 3' v3 from Ma2019.* ***(B)*** *The number of UMIs mapped to Merkel polyomavirus from two samples from patient 2586 (pre and post-immunotherapy treatment) sequenced using 10x 3' v2 and one sample from patient 9245 (post-immunotherapy treatment) sequenced using 10x 5' from Paulson2018.* ***(C)*** *The number of Fusobacterium reads per cell across different exposure conditions (regardless of cell type) from Robinson2023-plexWell.* ***(D)*** *The percentage of cells infected by Fusobacterium per cell type grouped by condition from Robinson2023-10x.* ***(E)*** *The percentage of cells infected by Hepatitis B per cell type (tumor vs. non-tumor) from sample H73a (sequenced using 3' v3) from Ma2019.* ***(F)*** *The percentage of cells infected by Merkel polyomavirus per cell type (tumor vs. non-tumor) from patient 9245 (sequenced using 5').* * p-value < 0.05; ** p-value < 0.01; *** p-value < 0.001.

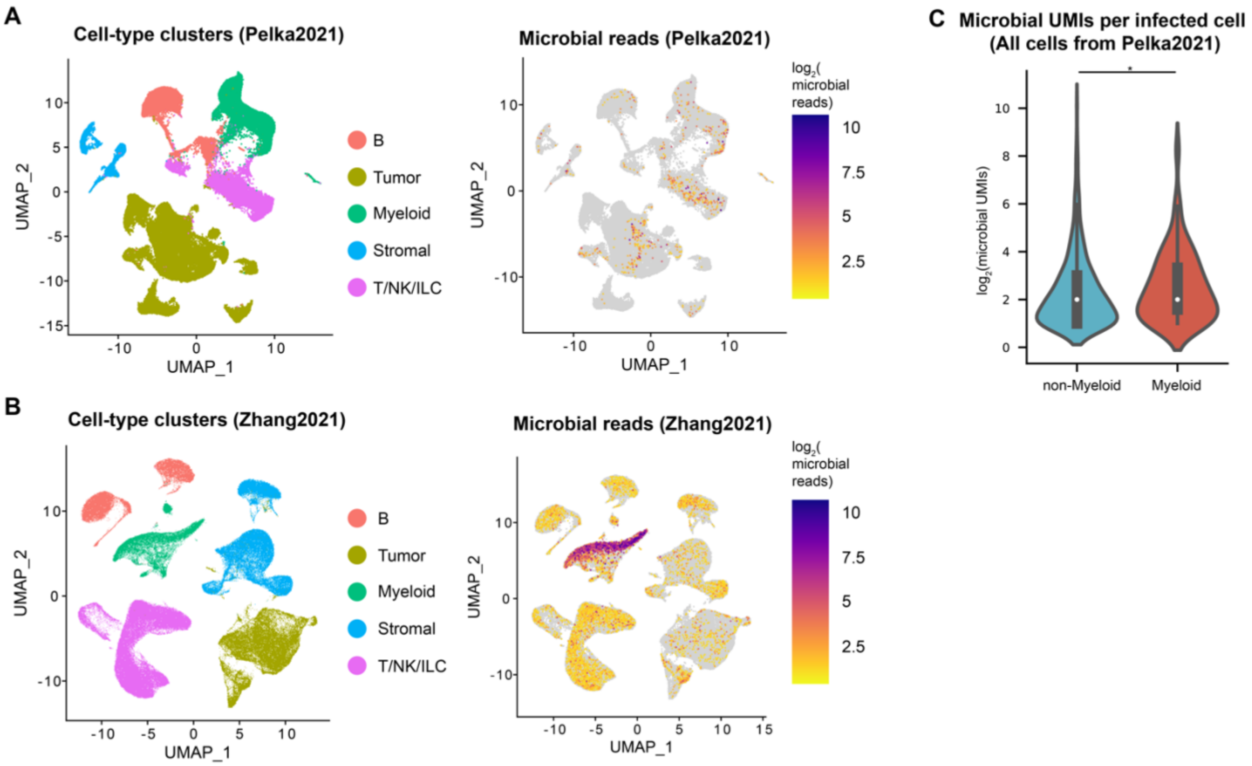


***Figure S3: Extended analysis of patient scRNA-seq datasets (Pelka2021 and Zhang2021)***

***(A)*** *UMAP analysis of cells analyzed from Pelka2021 colored by the cell type annotations (determined by the original authors) (left) and the number of microbial reads identified by CSI-Microbes.* ***(B)*** *UMAP analysis of cells analyzed from Zhang2021 colored by the cell type annotations (determined by the original authors) (left) and the number of microbial reads identified by CSI-Microbes (right).* ***(C)*** *The number of microbial UMIs per infected cells divided into myeloid and non-myeloid cells from Pelka2021.* * p-value < 0.05; ** p-value < 0.01; *** p-value < 0.001.

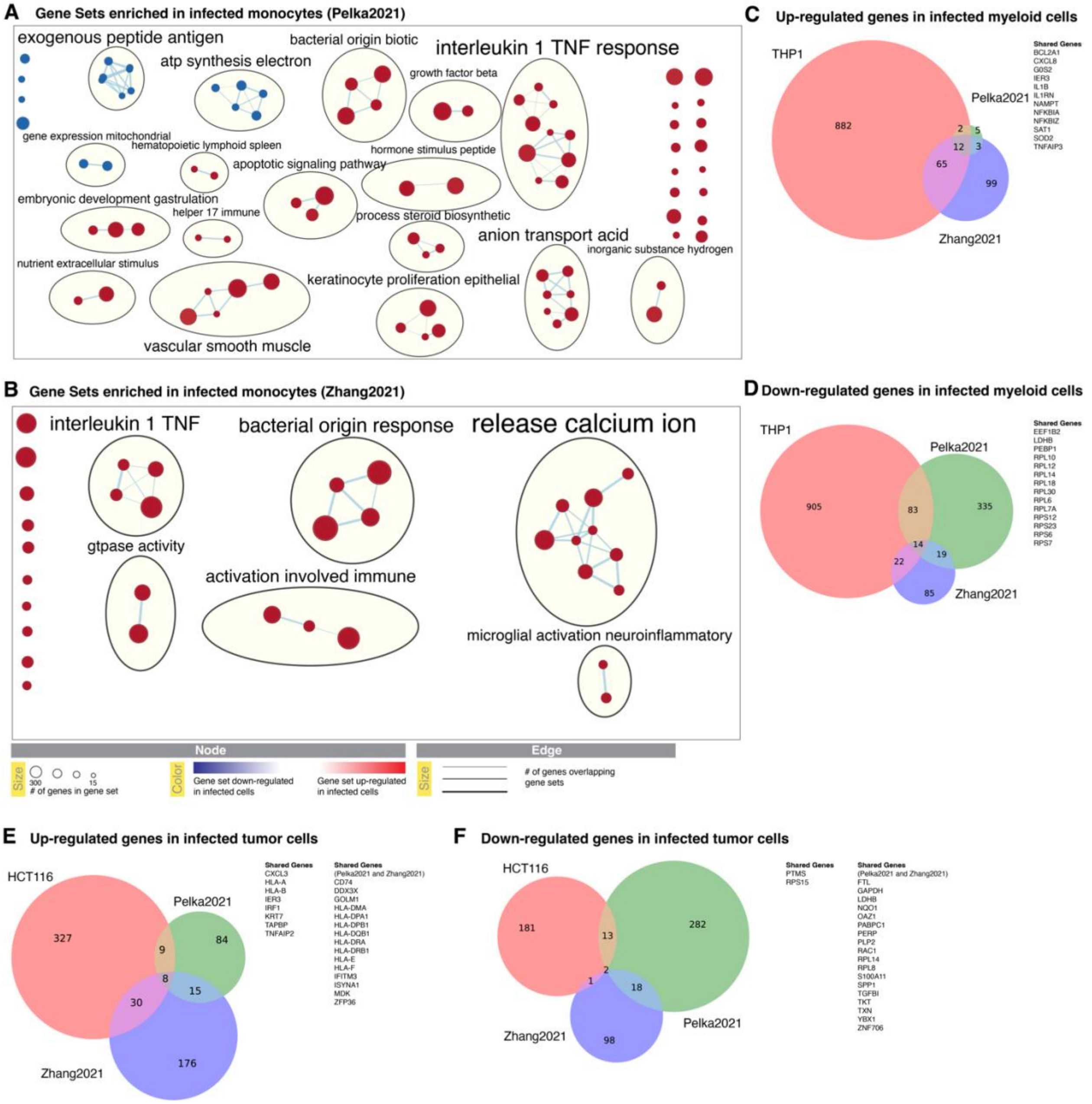


***Figure S4: Extended analysis of host transcriptomic changes associated with bacterial infection.***

***(A)*** *Annotated clusters of gene ontology (GO) biological processes (BP) gene sets up-regulated (red) or down-regulated (blue) in infected monocytes in Pelka2021.* ***(B)*** *Annotated clusters of GO BP gene sets up-regulated (red) or down-regulated (blue) in infected monocytes in Zhang2021.* ***(C)*** *The overlap of individual differentially expressed genes (DEGs) up regulated in infected myeloid cells in Pelka2021, infected myeloid cells in Zhang2021 and THP1 cells exposed to live F. nucleatum (compared to unexposed THP1 cells).* ***(D)*** *The overlap of down-regulated DEGs in infected myeloid cells in Pelka2021, infected myeloid cells in Zhang2021 and THP1 cells exposed* *to live F. nucleatum (compared to unexposed THP1 cells).* ***(E)*** *The overlap of up-regulated DEGs in infected tumor cells in Pelka2021, infected tumor cells in Zhang2021 and HCT116 cells exposed to live F. nucleatum (compared to unexposed HCT116 cells).* ***(F)*** *The overlap of down-regulated DEGs in infected tumor cells in Pelka2021, infected tumor cells in Zhang2021 and HCT116 cells exposed to live F. nucleatum (compared to unexposed HCT116 cells).*
